# Signatures of selection in pleiotropic genes involved in insect neuronal and immune systems

**DOI:** 10.64898/2026.03.02.708257

**Authors:** Sowmya Senthilkumar, Reese A. Martin, Ann T. Tate

**Affiliations:** Department of Biological Sciences, Vanderbilt University, Nashville, TN, USA; Evolutionary Studies Initiative, Vanderbilt University, Nashville, TN, USA

**Keywords:** neuroimmunology, neurodegenerative, evolvability, adaptation, evolutionary statistics Word Count: 4443

## Abstract

Pleiotropy, where a single gene contributes to multiple biological functions, plays a central role in shaping evolutionary constraints. The nervous and immune systems are tightly integrated both functionally and genetically, but it is not clear whether pleiotropy constrains adaptation in each system or contributes to disease. Building on evidence that genes that are pleiotropic between development and immunity in *Drosophila melanogaster* evolve slower than immune-specific genes, we tested whether pleiotropic neuro-immune genes likewise exhibit reduced evolutionary rates compared to genes functioning solely in neuronal or immune processes, and whether such evolutionary constraint is associated with human neurological disease. We identified immune, neuronal, and neuro-immune genes in *D. melanogaster* using Gene Ontology annotations, estimated their evolutionary rates across 12 *Drosophila* species, and investigated the breadth of their expression across developmental stages as a complementary proxy for pleiotropy. We further curated associations between the human orthologs for these genes and neurological disease and tested whether their evolutionary rates and pleiotropic status predicted disease involvement. We found that pleiotropic genes exhibited significantly lower dN/dS values and were less stage-specific than non-pleiotropic immune genes. Slower-evolving genes were more likely to be associated with human neurological diseases but this pattern was strongest for non-pleiotropic neuronal genes, suggesting that pleiotropy alone is not the strongest predictor of disease. Our results therefore indicate that dN/dS could be a stronger predictor of disease association than pleiotropy, raising new questions about the maintenance of pleiotropy in evolutionarily dynamic physiological systems.

## Introduction

Pleiotropy is the phenomenon where one gene affects multiple traits, and it plays an important role in shaping evolutionary trajectories (Fraïsse et al., 2018). Because mutations in pleiotropic genes can alter functions across a number of physiological systems, they are frequently subject to purifying selection, constraining their evolutionary potential (Fraïsse et al., 2018). This trade-off between multiplicity of function and evolability raises questions about how pleiotropic genes evolve when exposed to conflicting selective pressures.

As with most physiological systems that have distinct developmental trajectories and functions, nervous and immune systems have traditionally been studied independently from one another. However, growing evidence hints at substantial molecular and cellular interplay between these systems. For example, the survival of *Xenopus* embryos decreased significantly when injected with bacteria if they lacked brains (Herrera-Rincon, C, et al. 2020), revealing that very early brain development plays a central role in innate immunity (Dantzer, 2018; Kraus, 2021).

Neurons release immunomodulatory neurotransmitters that can either amplify or suppress immune responses, while immune cells produce neuroactive cytokines that influence neuronal function (Kipnis, 2018). West Nile virus, for example, disrupts the balance between the immune and nervous systems by activating genes with multiple roles in each system, suggesting that viral infections may impair neurotransmission by triggering immune responses that negatively affect neuronal function (Maximova 2021). Additionally, both the nervous and immune systems undergo extensive developmental regulation, and genes expressed across multiple life stages often experience strong evolutionary constraint (Liao and Zhang, 2006).

This functional overlap poses interesting questions about the evolutionary dynamics of pleiotropic immune and neuronal genes. After all, the immune system is under pressure to rapidly adapt to pathogens, and many immune genes show elevated levels of positive or balancing selection (Sackton et al., 2007, Williams, 2023). While some neuronal pathways show lineage-specific diversification consistent with adaptation (Tuller 2008, Yin 2015), many genes expressed in neural tissues evolve under strong purifying selection (Mccoy, 2023). For example, arylsulfatase A (ARSA) and galactocerebrosidase (GALC), lysosomal enzymes required for the metabolism of a major lipid component of the myelin sheath that insulates axons and enables rapid nerve impulse transmission, are highly conserved across vertebrates and exhibit low dN/dS ratios, consistent with strong purifying selection on genes essential for white-matter function (Luetzen, et al., 2025). Recent work suggests that immune genes with developmental roles evolve more slowly than those dedicated solely to immunity (Williams, 2023), raising the possibility that neuro-immune pleiotropy similarly constrains immune gene evolution. This idea is supported by the fact that many neuronal genes have critical developmental functions, suggesting they may be constrained similarly to developmental genes. If true, this would suggest that the adaptive potential of immune defenses is not solely dictated by pathogen pressure but also by the evolutionary constraints imposed by their additional roles in neural processes.

The relevance of this pleiotropic conflict extends beyond basic biology into the clinic. Many genes associated with neurological diseases such as Alzheimer’s and Parkinson’s also participate in immune defense (Provenzano et al., 2021; Bellou et al., 2020). For example, a patient-derived BCL11B mutation (N441K) causes combined immune and neurological defects. In mice, the equivalent mutation (N440K) led to both abnormal NK/ILC1-like immune cell development in the thymus and reduced cortical neurons (Okuyama, K, 2023). Hyperactive immune responses, moreover, protect against infection but can trigger neuroinflammation, accelerating neurodegeneration (Provenzano et al., 2021). Understanding how the shared genes and pathways navigate competing pressures could reveal why certain neurological disorders persist in populations despite their costs to health and fitness.

Many of the mechanisms controlling neural development and immune activation are conserved between insects and mammals (Lye, 2018). The extensive genetic and genomic data and tools available for the fruit fly *Drosophila melanogaster* make it an excellent model organism to explore how pleiotropy influences selection pressures and organismal phenotypes associated with infection and neurodegenerative diseases (Lye, 2018). By determining the evolutionary rates (dN/dS) of pleiotropic immune-neuronal genes across 12 *D. melanogaster* species, we tested whether nervous system functions impose evolutionary constraints on immune adaptation. We predicted that immune genes with neuronal roles would have lower dN/dS values than immune-specific genes, reflecting stronger purifying selection. We further predicted that pleiotropic genes would have more associations with neurological disorders compared to non-pleiotropic genes after controlling for evolutionary rate, as their constrained evolution may reflect strict roles in neuronal development and maintenance. Support for these predictions would shed light on how pleiotropy shapes the molecular evolution of interconnected biological systems more broadly as well as trade-offs between immunity and nervous system maintenance.

## Methods

### Categorization of non-pleiotropic and pleiotropic immune and neuronal genes

We extracted immune-related and neuronal gene lists for *D. melanogaster* from FlyBase by manually collecting GO-annotated genes tagged with the “immune system process” (GO: 0002376) or “neuron development” (GO: 0048666) terms respectively. From these, we identified 724 neuronal genes, 584 immune genes, and 96 genes with overlapping functions, which we defined as pleiotropic. In this study, genes binned in a “non-pleiotropic” class does not mean they have *no* pleiotropic function; after all, most genes are likely pleiotropic if enough traits are considered (Boyle, 2017); these genes simply do not exhibit pleiotropy between our two major systems of interest (see additional discussion of this approach in Willams et al 2023). The full list of genes in each group is included in Supplementary Table 1.

### Compiling sequences for analysis

We followed the protocol established in Williams *et al* 2023, where a subset of the *Drosophila* clade was used to determine the rate of evolution of genes that were pleiotropic between developmental and immune processes. First, we obtained coding sequences (CDSs) for *Drosophila melanogaster* using the FlyBase Sequence Downloader tool with FlyBase gene IDs. From an archived FlyBase version (FB2017_05), we retrieved ortholog lists for 12 Drosophila species (*D. ananassae, D. erecta, D. grimshawi, D. mojavensis, D. persimilis, D. pseudoobscura, D. sechellia, D. simulans, D. virilis, D. willistoni, and D. yakuba*). The previous study (Williams *et al*. 2023) demonstrated that the main results were relatively insensitive to the use of this broader species set versus smaller subgroups despite the risk of saturated dS, so we chose to repeat the 12 species approach here for consistency. We downloaded CDSs for each gene of interest across species using the archived FlyBase Sequence Downloader tool. Using scripts adapted from the supplementary materials of that study, we parsed FlyBase sequence IDs for each species.

We aligned sequences in codon space and trimmed them using Gblocks v0.91b (Castresana 2000). When we encountered paralogs, we excluded those genes from alignment. We retained only genes with ≥2 high-quality orthologous CDSs for downstream analysis, resulting in 545 neuronal, 428 immune, and 77 pleiotropic curated alignments. We used these filtered gene sets (rather than the initial GO-derived lists) for all evolutionary and biological-process count analyses described below.

### Biological counts as a measure of pleiotropy

To characterize the prevalence of pleiotropy within the filtered gene sets, we quantified the number of unique Gene Ontology (GO) biological process annotations per gene. For each gene included in the evolutionary analyses, GO analysis for each gene were retrieved using the FlyBase “ID Validator” tool, and biological process annotations were extracted from the resulting hit lists. The number of biological processes for each gene were then summarized for each functional category (neuronal, immune, pleiotropic; Supplemental Table 2). The biological process counts for each gene ID are listed in Supplemental Table 3.

### Calculating dN/dS values using PAML

The dN/dS ratio, which compares nonsynonymous to synonymous substitutions, served as our measure of evolutionary rate. A dN/dS > 1 indicates positive selection driving amino acid changes, while a lower ratio reflects purifying selection preserving protein structure (Jackowiak, 2014). Although values exceeding 1 are uncommon, relatively higher dN/dS ratios still suggest that a gene is experiencing stronger positive selection, or at least relaxed purifying selection, compared to others. We used these sequence files to calculate evolutionary rate measurements by running them through codeml site model M0 in PAML (v 4.10.7). Our PAML analysis successfully processed all gene lists (539 neuronal genes, 420 immune genes, and 74 pleiotropic genes). In addition to per-gene analyses, we generated concatenated alignments for each gene category to estimate overall category-level dN/dS values using PAML model M0. Finally, we performed Kruskal-Wallis tests followed by post hoc Dunn tests in R (v2026.01.0) to compare dN/dS values. The dN/dS values for each gene in each category are listed in Supplemental Table 4.

### RNA-seq expression data acquisition and preprocessing

RNA-seq gene expression data were obtained from the modENCODE developmental expression datasets curated in FlyBase. Specifically, we downloaded RPKM (Reads Per Kilobase per Million mapped reads) values from the FlyBase gene_rpkm_report_fb_*.tsv.gz files, which compile normalized RNA-seq expression estimates across developmental stages for *Drosophila melanogaster* genes. For each gene class (pleiotropic, non-pleiotropic neuronal, and non-pleiotropic immune), we filtered the full FlyBase expression matrix to retain only genes belonging to the corresponding category (Supplemental Tables 5-8). Gene identifiers were matched using FlyBase gene IDs (FBgn). Expression values were extracted for three representative developmental stages: larval stage 1 (L1), pupal stage 5 (P5), and adult male one day post-eclosion (AdM_Ecl_1days).

Embryonic stages were excluded from the main analysis because they are represented by multiple short time windows (e.g., 0–2 hr, 2–4 hr, etc.), which would disproportionately weight embryonic expression and artificially depress τ values when aggregated into a single developmental category. Restricting the analysis to larval, pupal, and adult stages allowed for balanced comparison of expression specificity across major post-embryonic developmental transitions, although results from including the embryonic stage are represented in the Supplemental Materials.

### Calculation of gene expression specificity (τ)

To quantify developmental expression specificity, we calculated the τ (tau) metric as originally described by Yanai et al. (2005). Tau provides a normalized measure of how restricted a gene’s expression is across tissues or developmental stages, ranging from 0 (broad expression across all stages) to 1 (highly stage-specific expression).

For each gene, τ was calculated across the three developmental stages using the following equation:

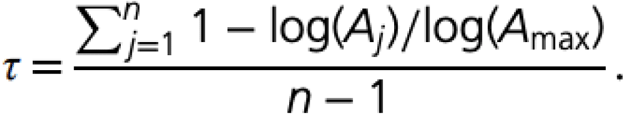

where *n* represents the number of stages, Aⱼ the expression level (RPKM) of the gene at stage/tissue j, and A_max_ is the maximum expression value of that genes across all stages. Prior to log transformation, a pseudocount of 1 was added to all RPKM values to avoid undefined logarithms for genes with zero expression at one or more stages. Tau values were computed independently for each gene, resulting in a gene-level distribution of τ values for each gene class. Differences in τ distributions among pleiotropic, non-pleiotropic neuronal, and non-pleiotropic immune gene classes were assessed using a Kruskal-Wallis rank-sum test.

When the Kruskal–Wallis test indicated a significant overall effect, post hoc pairwise comparisons between gene classes were performed using Dunn’s test. Resulting p-values were adjusted for multiple comparisons using the Benjamini-Hochberg false discovery rate (FDR) correction. The specific tau values for each gene is listed in Supplemental Table 9.

### Gene ontology cellular component enrichment analysis

Gene Ontology (GO) enrichment analyses were performed using PANTHER, focusing specifically on the Cellular Component (GO-CC) ontology. Enrichment results were reported as fold enrichment values, representing the ratio of observed gene counts in a given GO category relative to the expected number based on background frequency. P values were corrected for multiple testing using false discovery rate (FDR) correction as implemented by PANTHER. Fold enrichment values were log₂ transformed and log₂ fold enrichment = 0 corresponds to a fold enrichment of 1. To facilitate comparison across gene categories while minimizing visual clutter, a shared set of GO Cellular Component terms was selected for plotting. Specifically, for each gene category, the top N GO terms (N = 10) were ranked by FDR-adjusted p value, prioritizing the most statistically significant enrichments. The union of these top-ranked terms across the three gene categories was used to define the final set of GO terms displayed, resulting in approximately 22 GO terms after accounting for overlap among categories. Complete enrichment statistics, including GO identifiers, gene counts, and FDR values, are provided in the supplementary tables 10-12.

### The relationship between dN/dS and neuronal diseases

To examine whether dN/dS was associated with neuronal disease, we annotated each gene in our dataset for known human neurological disease associations. We used the FlyBase ID Validator tool to retrieve human ortholog information and associated Human Disease Ontology annotations for each *D. melanogaster* gene. For every gene, we manually screened all annotations for relevance to neurological or neurodevelopmental disease categories, using keywords including but not limited to: *neurodevelopmental disorder*, *neurodegenerative disease*, *motor neuron disease*, *Parkinson disease*, *Alzheimer disease*, *ataxia*, and *intellectual disability*. Based on these annotations, each gene was assigned a binary neuronal-disease variable, where 1 indicated at least one associated neurological disease term in its human ortholog(s), and 0 indicated the absence of any such disease annotation.

To examine the relationship between evolutionary rate and neuronal disease association, we fit a binomial logistic regression model with neuronal disease status as the response variable and both gene class (pleiotropic vs. non-pleiotropic) and gene-specific dN/dS as predictors. Models were fit in R using the glm function with a binomial distribution. Test statistics and associated p-values were obtained from the resulting analysis.

## Results

### Pleiotropic genes are involved in more biological processes than non-pleiotropic immune and neuronal genes

To determine if the gene we identified as pleiotropic between immunity and the nervous system were in general more pleiotropic than purely immune or nervous genes, we counted the total number of unique biological processes associated with each gene in our study (Fig. S1). We found that genes that were pleiotropic between immunity and the nervous system are associated with significantly more biological processes than non-pleiotropic immune (P.adj = <0.0001) and neuronal (P.adj < 0.0001) genes (Fig. 1). There was no significant difference in biological process counts between the non-pleiotropic groups (P.adj = .06).

**Figure 1.**
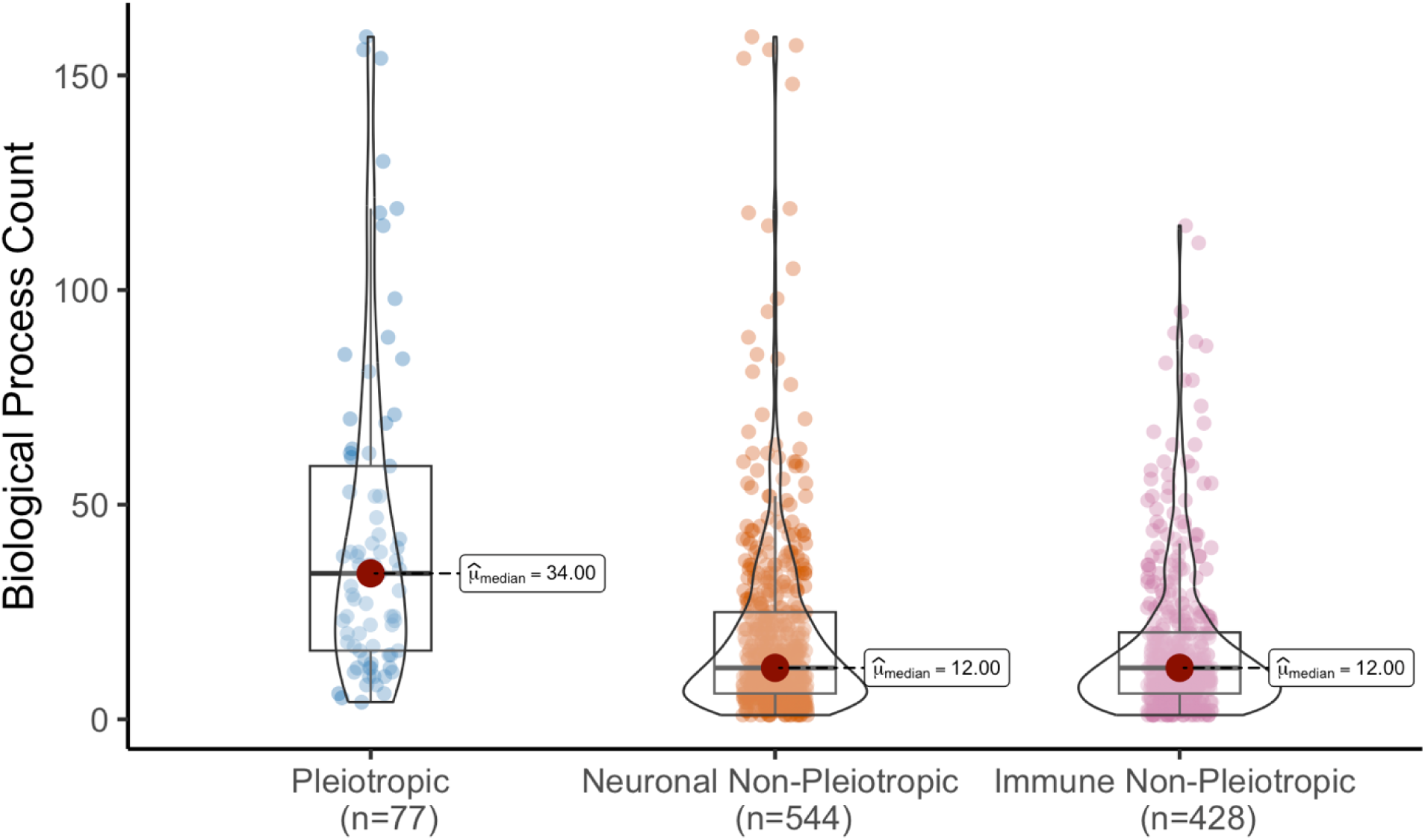
Number of biological processes associated with each class of genes. Each gene was categorized into a class based on its annotated function(s). The number of unique annotated Gene Ontology (GO) biological processes was determined for each gene. These counts reflect the number of functional roles a gene plays in various biological processes and are shown on the y-axis. The violin plots show the distribution density of biological process counts across genes in each category. Individual points represent individual genes. Post hoc pairwise Dunn tests were used to calculate pairwise comparisons, and p values were adjusted for false discovery rate (Benjamini-Hochberg FDR).

### Pleiotropic and non-pleiotropic gene classes show distinct patterns of cellular component enrichment

We performed Gene Ontology (GO) Cellular Component enrichment analyses on pleiotropic, non-pleiotropic neuronal, and non-pleiotropic immune gene sets and compared patterns of enrichment across gene classes (Figure 2). Across the top significantly enriched GO cellular component terms for each gene set, pleiotropic genes showed enrichment for a wide range of broadly defined cellular structures, including *cell body*, *cell projection*, *cytoplasmic vesicle*, *intracellular vesicle*, and *plasma membrane-bounded cell projection* (Figure 2). In contrast, non-pleiotropic neuronal genes exhibited stronger enrichment for neuron-specific compartments such as *axon*, *somatodendritic compartment*, *neuron projection*, and *neuronal cell body*, consistent with functional specialization within the nervous system. Non-pleiotropic immune genes were enriched for a distinct set of cellular components, including *extracellular region*, *extracellular space*, *perivitelline space*, and immune-related complexes such as the *calcineurin complex*, indicating specialization toward immune-associated cellular contexts.

**Figure 2:**
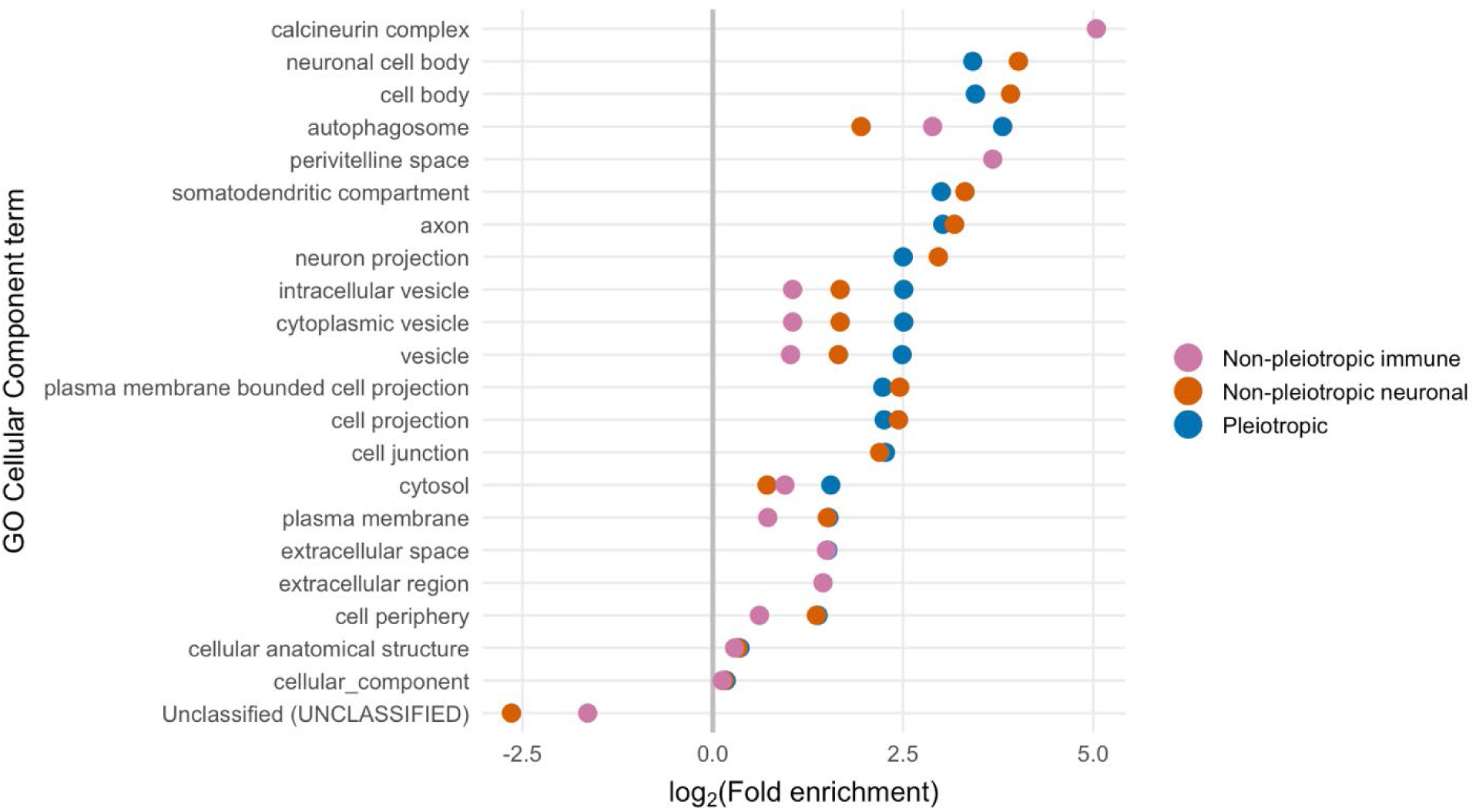
Gene Ontology (GO) Cellular Component enrichment analysis comparing pleiotropic, non-pleiotropic neuronal, and non-pleiotropic immune gene sets. The dot plot displays the log₂ fold enrichment for significantly enriched GO cellular component terms. Each row corresponds to a GO Cellular Component term, and each dot represents a statistically significant enrichment for a given gene category (FDR ≤ 0.05). Dot color denotes gene category. Only GO terms that were among the top FDR-significant terms across gene categories are shown. Absence of a dot indicates that the GO term was not significantly enriched in that gene category.

Several GO terms were shared across gene classes but differed in the magnitude of enrichment, with pleiotropic genes generally showing moderate enrichment across many cellular components, whereas non-pleiotropic gene classes displayed stronger enrichment in fewer, more specialized compartments.

### Pleiotropic genes evolve more slowly than non-pleiotropic immune and neuronal genes

To measure the evolutionary rate of pleiotropic genes compared to the non-pleiotropic genes, we ran PAML model M0 with concatenated lists of pleiotropic and non-pleiotropic genes. The PAML model on the gene lists gave median dN/dS of .0516, .0592, and .0748 for pleiotropic genes, neuronal genes, and immune genes respectively.

Pleiotropic genes evolve significantly more slowly than non-pleiotropic immune genes (P.adj = 7.87e-04; Figure 3). There was no significant difference between the pleiotropic genes and non-pleiotropic neuronal genes (P.adj= .29), but non-pleiotropic immune genes did have significantly higher dN/dS values than non-pleiotropic neuronal genes (P.adj = 8.19e-06).

**Figure 3.**
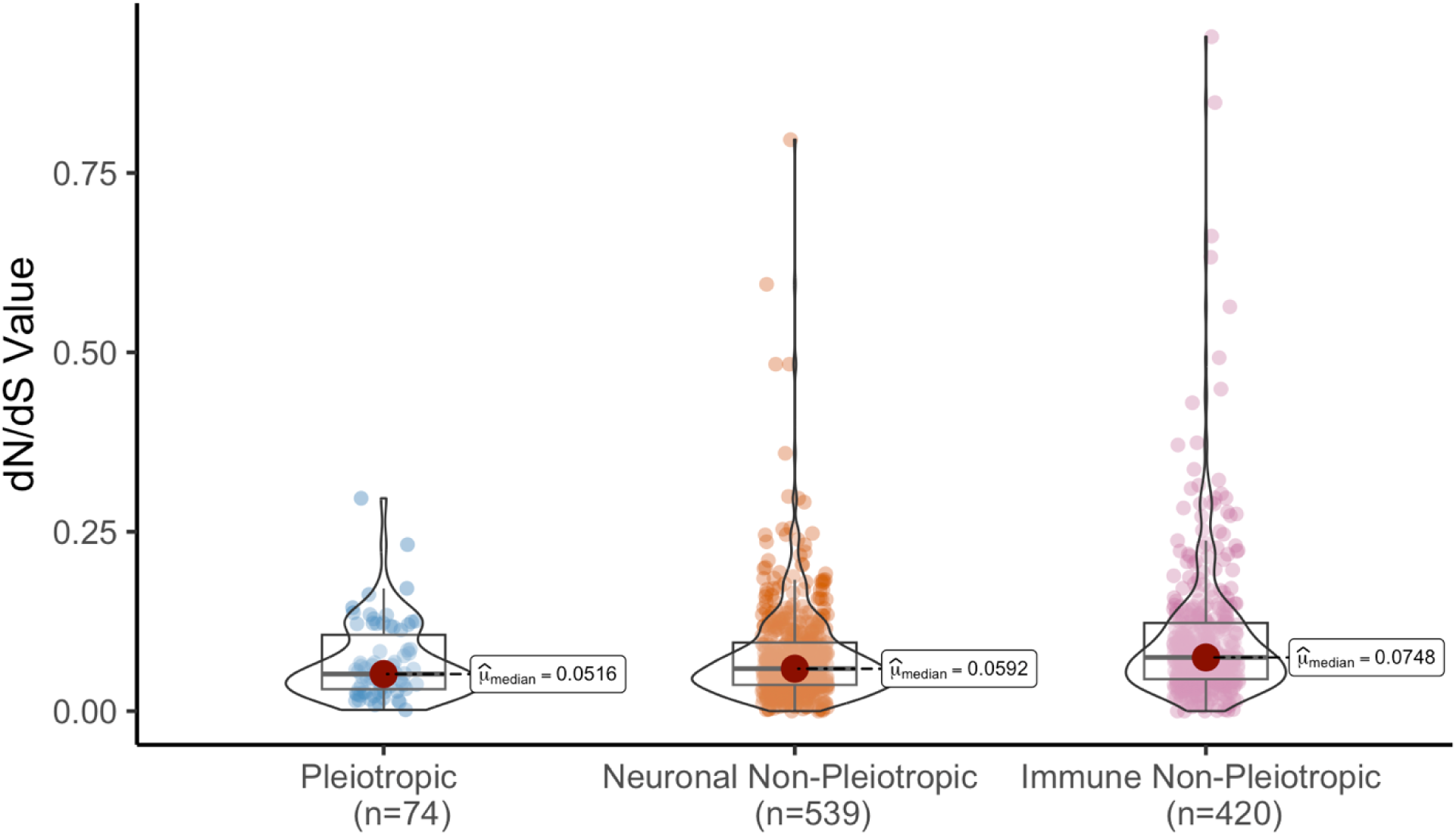
Violin plot showing the distribution of evolutionary rates, as measured by nonsynonymous to synonymous substitution ratios (dN/dS), for three gene categories: pleiotropic, nonpleiotropic neuronal, and nonpleiotropic immune. Each violin plot represents the distribution of dN/dS values for genes in a given category. The width of the violin corresponds to the density of genes at different dN/dS values. Higher dN/dS values suggesting positive or relaxed purifying selection and lower dN/dS values suggest stronger purifying selection. Post hoc pairwise Dunn tests were used to calculate pairwise comparisons, and p values were adjusted for false discovery rate (Benjamini-Hochberg FDR).

### Pleiotropic genes are more broadly expressed across life stages than non-pleiotropic ones

Pleiotropic genes are hypothesized to be more broadly expressed across stages than non-pleiotropic immune genes, reflecting their role in multiple or housekeeping functions throughout life (Yanai et al. 2005; Williams, 2023). To test this hypothesis, we assessed the uniformity of gene expression across larval, pupal and adult stages in *D. melanogaster* by comparing the distribution of tau values (τ) for each gene in each gene class (Fig. S2). Lower τ values indicate broader expression across life stages. Genes in the non-pleiotropic immune class were significantly more stage specific than non-pleiotropic neuronal genes (P.adj = 2.96e-03) and pleiotropic genes (P.adj = .04; Figure 4). Pleiotropic genes and non-pleiotropic neuronal genes had similar stage specificity values (P.adj=0.59; Figure 4). All significant effects disappeared when the embryonic stage was added to the analysis, due to an extreme excess in the expression of embryo-only genes (Fig. S3).

**Figure 4.**
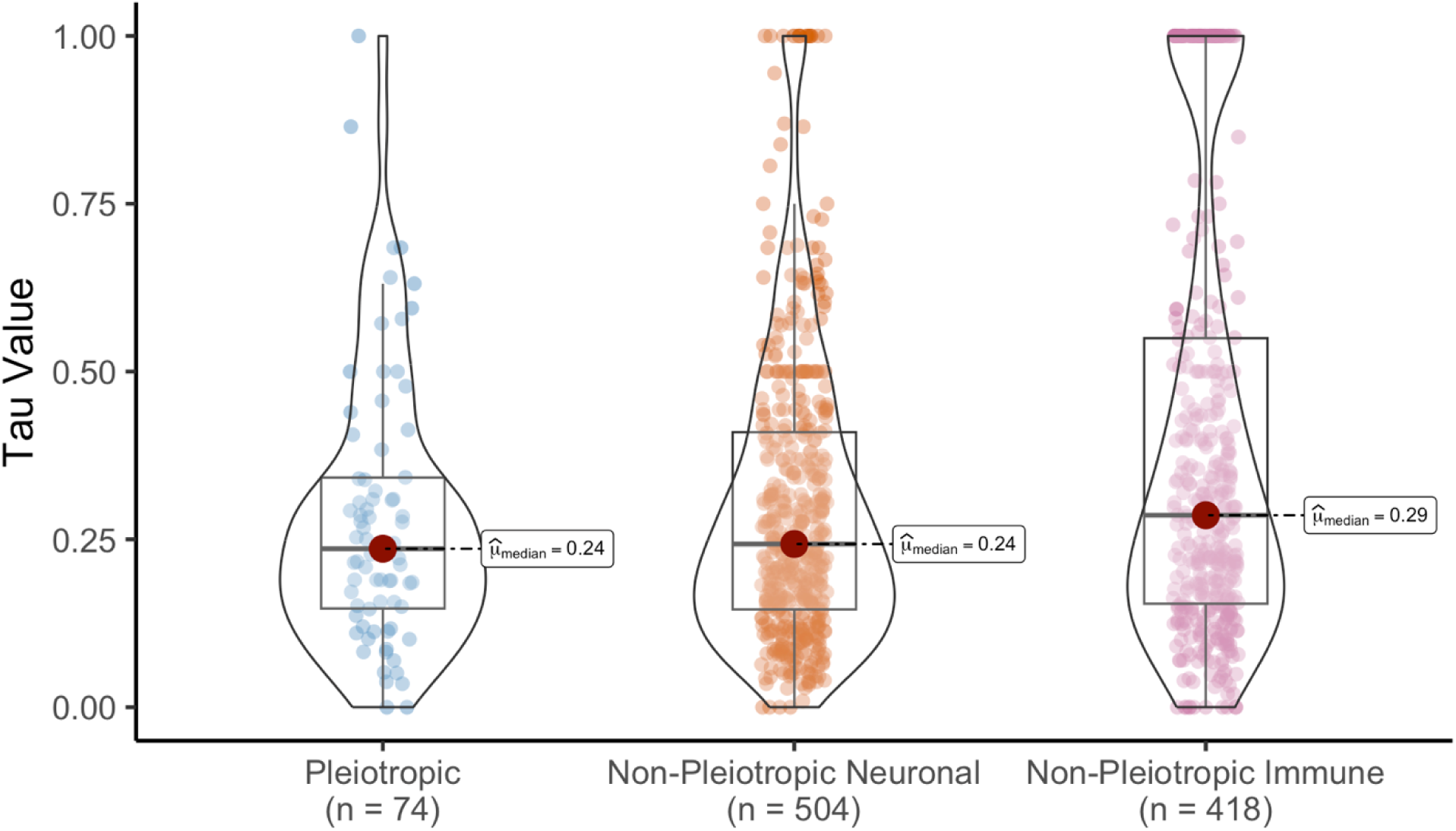
Gene expression specificity (tau) compared across larval, pupal, and adult stages for pleiotropic and non-pleiotropic gene classes. Tau values represent the gene expression specificity and range from 0 (broadly expressed across stages) to 1 (specific to a stage). Each violin represents the density distribution of tau values for a gene category. Individual dots represent tau values for each gene within the category. Median tau values are labeled for each group. P values were statistically obtained by the Kruskal-Wallis Test. Post hoc pairwise Dunn tests were used to calculate pairwise comparisons, and p values were adjusted for false discovery rate (Benjamini-Hochberg FDR).

### Genes under stronger purifying selection are more strongly associated with human neuronal diseases

To examine the relationship between evolutionary rate and neuronal disease association, we performed a binomial logistic regression with neuronal disease status as the response variable and both dN/dS and gene class (pleiotropic vs. non-pleiotropic) as predictors (Figure 5). Across all genes, dN/dS was a significant predictor of disease association. Specifically, dN/dS showed a strong negative relationship with the probability of neuronal disease involvement (Supplmental Table 13; coefficient estimate = -13.36, SE = 3.61, p = 0.0002), indicating that genes evolving under stronger purifying selection were more likely to be associated with neurological disease.

**Figure 5.**
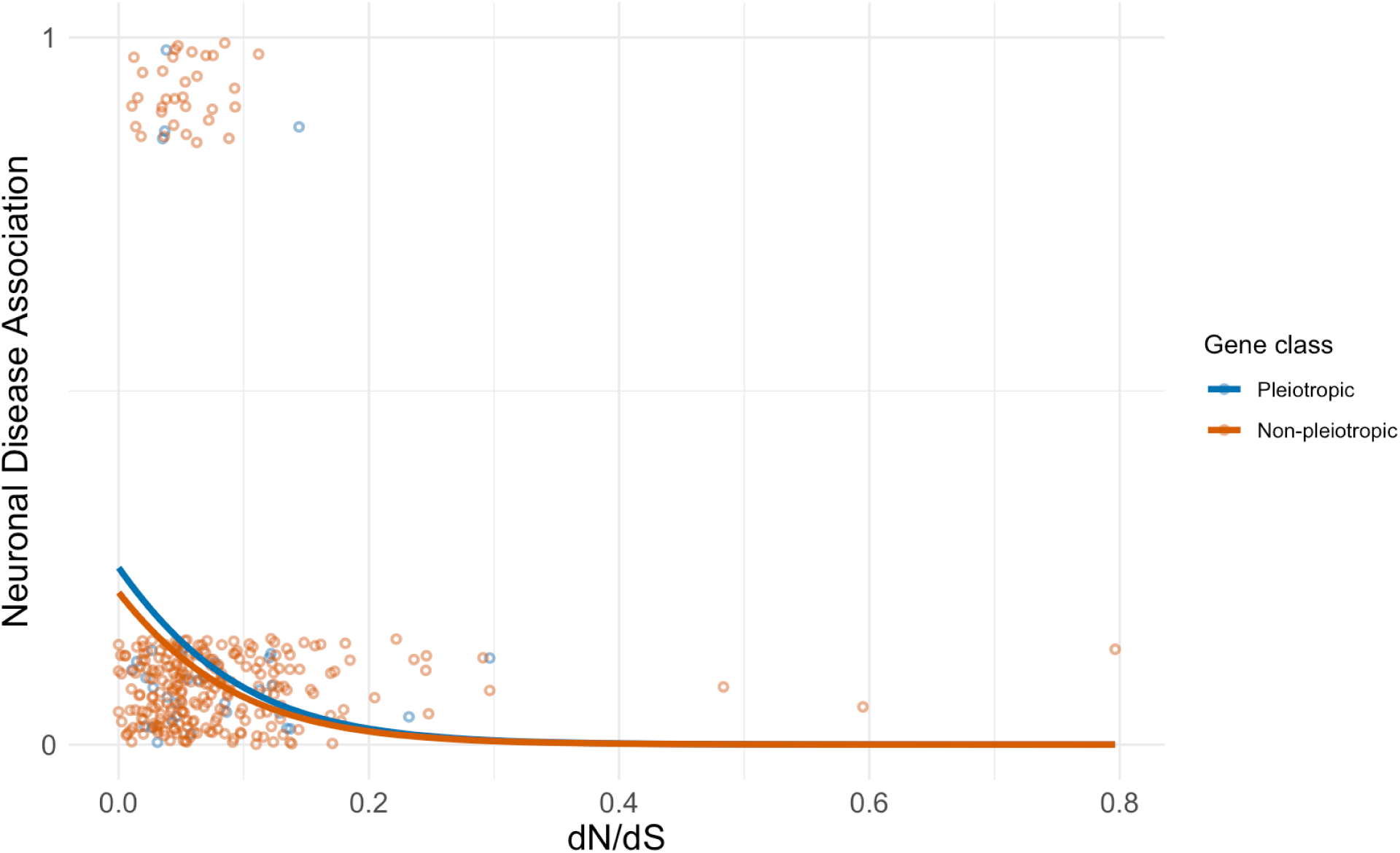
Binomial distribution showing the relationship between rates of evolution (dN/dS values) and neuronal disease association for individual genes in pleiotropic and non-pleiotropic gene classes. Logistic regression curves are shown for pleiotropic genes (blue) and non-pleiotropic genes (orange). The x-axis represents dN/dS values, and the y-axis represents the binary outcome of neuronal disease association (0 = no association, 1 = associated), as determined from human disease annotations linked to each gene’s human ortholog in FlyBase. Each point represents an individual gene, and the fitted curves show the predicted probability of neuronal disease association across increasing dN/dS values for each gene class. All dots are either 0 or 1 on the y axis but are jittered for ease of viewing.

In contrast, gene class was not a significant predictor after accounting for evolutionary rate (coefficient estimate = - 0.20, SE = 0.37, z = -0.53, p = 0.60). The fitted logistic curves for pleiotropic and non-pleiotropic genes show similar trends, with both gene classes exhibiting decreasing probabilities of disease association as dN/dS increases.

## Discussion

In this study we tested the hypothesis that pleiotropy constrains the evolution of genes involved in both immune and neuronal systems relative to genes that function in only one of the two processes. Our results suggest that pleiotropic genes do evolve more slowly than non-pleiotropic immune genes but evolve at a similar rate compared to non-pleiotropic neuronal genes (Figure 3). Our results further suggest that neurological disease associations are more strongly linked to genes evolving under stronger purifying selection, and that this relationship is better explained by evolutionary constraint than by pleiotropy alone.

The pleiotropic genes in our study were associated with significantly more GO biological processes than either of the non-pleiotropic groups (Figure 1). This estimate, which includes all GO biological processes and not just neuronal and immune ones, provides complementary support that these genes are more pleiotropic overall than those in the non-pleiotropic categories. The highest median dN/dS value observed in this study was associated with the non-pleiotropic immune genes. This aligns with prior evidence that many immune genes, particularly those without pleiotropic constraints, undergo rapid adaptive evolution (Williams, 2023). The median dN/dS values for the non-pleiotropic neuronal and non-pleiotropic immune genes were statistically different from each other, suggesting that at least some neuronal non-pleiotropic genes evolve under less constraint than their pleiotropic counterparts. Previous work suggests that genes highly expressed in cortical regions of the brain have lower evolutionary rates than genes highly expressed in subcortical regions (Tuller 2008), providing a biological explanation for our observation of a large spread in the distribution of dN/dS values in non-pleiotropic neuronal genes (Figure 3). Consistent with this interpretation, non-pleiotropic neuronal genes also show strong enrichment in highly specialized neuronal compartments such as axons and somatic and dendritic regions (Figure 2), suggesting that cellular compartmentalization may further influence variation in evolutionary rate.

The statistical relationship between neuronal pleiotropy and evolutionary rate observed in this study builds directly on the developmental pleiotropy framework established by Williams (2023), which serves as the foundation for our analysis. Williams demonstrated that developmental genes, as well as genes with dual roles in immunity and development, evolve more slowly and experience stronger purifying selection than non-pleiotropic immune genes.

Consistent with this framework, just as developmental genes are predicted to be under purifying selection to ensure a faithful developmental program, the reduced evolutionary rate of pleiotropic neuronal genes in our study may arise from their conserved effects on neural network maintenance (Mccoy, 2023).

Notably, neuro-immune pleiotropic genes show evolutionary patterns that more closely resemble other neuronal genes than immune genes. The lower dN/dS values observed in pleiotropic neuronal genes are consistent with stronger purifying selection compared to non-pleiotropic immune genes (Figure 3). This may relate to functional constraints on immunity that protects delicate neural tissue from potentially damaging inflammation (Buckley, 2022).

Although once thought to lack immune function, neurons and glial cells do express many immune-related proteins in restricted spatial and temporal patterns consistent with non-immunological roles. For instance, major histocompatibility complex class I (MHCI) proteins, which are central to adaptive immunity, are also expressed in neurons, where they contribute to neural signaling and plasticity (Elmer et al., 2012). These findings suggests that immune-related genes expressed in neuronal functions may experience stronger purifying selection due to their dual roles in maintaining neural integrity. Consistent with this dual-use framework, pleiotropic genes in our dataset are enriched in neuronal compartments such as axons and focused neuronal regions (Figure 2), suggesting that immune-related genes co-opted for neuronal functions may experience stronger purifying selection due to their essential roles in maintaining neural integrity.

The observation that pleiotropic genes are more broadly expressed across developmental stages than non-pleiotropic immune genes is consistent with previous results on immune-developmental pleiotropy (Williams et al 2023) and provides insight into the roles these genes play throughout the organism’s development (Figure 2). Previous work has highlighted that genes involved in multiple biological processes often display conserved, widespread expression due to their essential roles in core cellular functions (Wagner, 2008). In *D. melanogaster*, for example, highly pleiotropic genes such as ELAV/Hu factors comprise a conserved family of RNA binding proteins (RBPs), maintain stable expression across neural lineages while also functioning in RNA processing, an interplay that restricts their evolutionary flexibility (Lee et al. 2021). Our observation that pleiotropic genes show similar expression breadth to non-pleiotropic neuronal genes raises additional questions about the underlying regulatory mechanisms that constrain or diversify gene expression. Neuronal genes are often broadly expressed during development due to the complexity of neural tissue formation and function (Crews, 2019). This similarity could suggest that neural development shares a demand for versatile gene expression, or alternatively, that some of the pleiotropic genes are particularly important for neurological functions throughout life. Notably, the non-pleiotropic immune gene set shows an enrichment of genes with τ values near 1, indicating a large subset of highly stage-specific immune genes. This pattern is consistent with immune genes that are tightly regulated and expressed predominantly during specific physiological windows, such as acute immune activation or pathogen exposure ((Lemaitre & Hoffmann, 2007; Buchon et al., 2014). In contrast, pleiotropic genes display fewer extreme τ values, suggesting broader expression across stages, likely reflecting their dual functional roles across systems. The abundance of highly stage-specific immune genes may reduce pleiotropic constraint within the immune-only gene set, allowing more specialized regulation (Alvarez-Ponce, 2017), whereas pleiotropic genes remain evolutionarily constrained due to their sustained expression and functional importance across both immune and neuronal contexts.

Genes with reduced evolutionary rates were more frequently associated with neuronal disease, regardless of whether they were pleiotropic or neuron-specific (Figure 5). This pattern suggests that strong purifying selection on genes involved in essential neural and immune-related processes, rather than pleiotropy alone, is associated with disease involvement. This was surprising because it contrasts to some extent with previous associations between pleiotropy and disease. For example, Ittisoponpisan et al. (2016) analyzed pleiotropic proteins as those in which variants in the same protein cause multiple distinct diseases and found that these proteins were more likely to be associated with disease, suggesting that genes with multiple functional roles are under stronger functional constraint. It is worth noting that our study links *Drosophila* evolutionary statistics to human ortholog-based disease annotations, and there is no guarantee that selection pressures across these very different taxa have remained consistent over time, despite reasonable conservation of immune and neuronal gene roles. Nevertheless, our results provide an evolutionary perspective that complements the network-based findings of previous work, suggesting that the apparent enrichment of disease associations in pleiotropic genes may be largely explained by their stronger evolutionary constraint rather than pleiotropy itself.

Further research with expanded datasets and disorder-specific analyses will help clarify how pleiotropy and evolutionary constraint interact to shape patterns of disease association.

Overall, our findings demonstrate that neuronal pleiotropy slows the evolution of immune genes in a manner similar to previously established constraints imposed by developmental pleiotropy. Neuro-immune pleiotropic genes in *D. melanogaster* evolve more slowly than immune-specific genes, are broadly expressed across life stages, and exhibit evolutionary patterns closely associated with neuronal genes. Importantly, we also show that *Drosophila* genes with human orthologs associated in neurological disorders tend to be those under the strongest purifying selection, underscoring that evolutionary conservation in flies can serve as a useful indicator of disease relevance in humans. Together, these patterns suggest that pleiotropy imposes strong functional constraints on essential neuro-immune genes, limiting their evolutionary flexibility and increasing vulnerability to deleterious mutations. Such constraints may help explain why variants in these genes are repeatedly implicated in neurological disorders, despite strong purifying selection. Future work integrating population genetic analyses, comparative genomics across additional taxa, and functional assays will help clarify the selective forces maintaining pleiotropy in immune genes and reveal why some highly constrained neuro-immune genes remain vulnerable points in the architecture of human neurological disease.

## Acknowledgements

We thank Allyson Ray and other members of the Tate lab for comments and discussion, and Alissa Williams for providing draft scripts and guidance. This work was supported by the National Institute of General Medical Sciences at the National Institutes of Health (grant number R35GM138007 to A.T.T.)

