## Supplemental Figures for "Signatures of selection in pleiotropic genes involved in insect neuronal and immune systems"

Senthilkumar, Martin, and Tate 2026

A

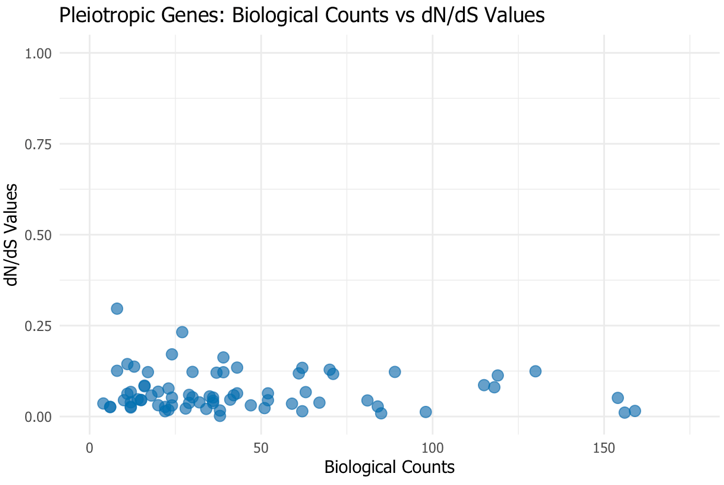

B

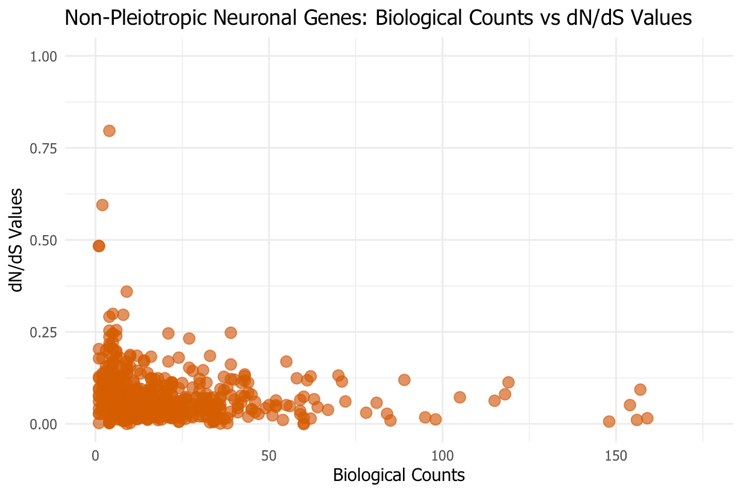

C

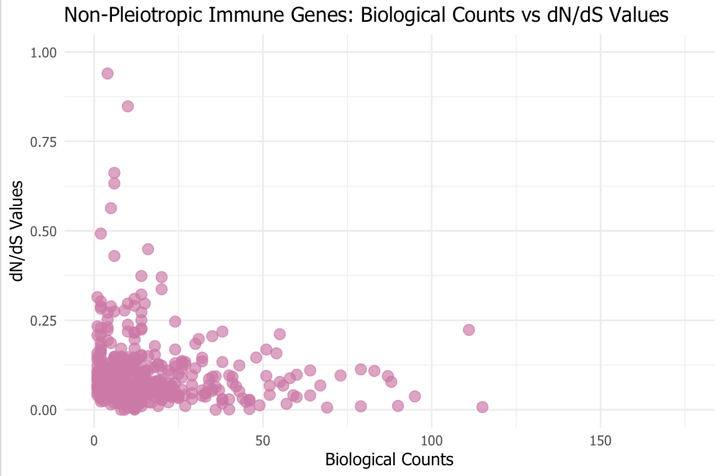

Supplemental Figure 1. Relationship between the number of biological counts and individual dN/dS values across gene categories. Scatter plots depict individual gene data points, illustrating the relationship between biological counts (x-axis) and dN/dS values (y-axis) for three gene groups: non-pleiotropic immune genes (A), non-pleiotropic neuronal genes (B), and pleiotropic genes (C). Each point represents a single gene, where biological counts indicate the number of biological processes associated with the gene, and dN/dS values reflect the ratio of nonsynonymous to synonymous substitutions, representing evolutionary selection pressure.

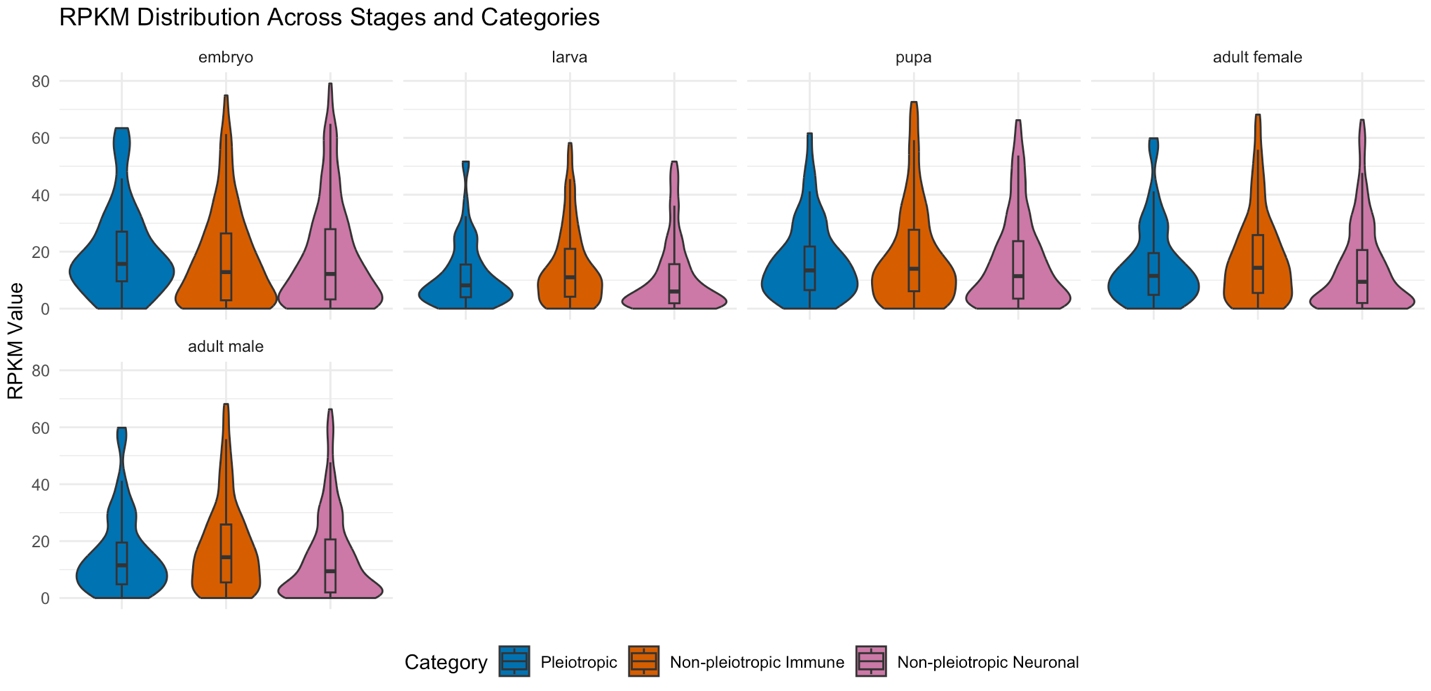

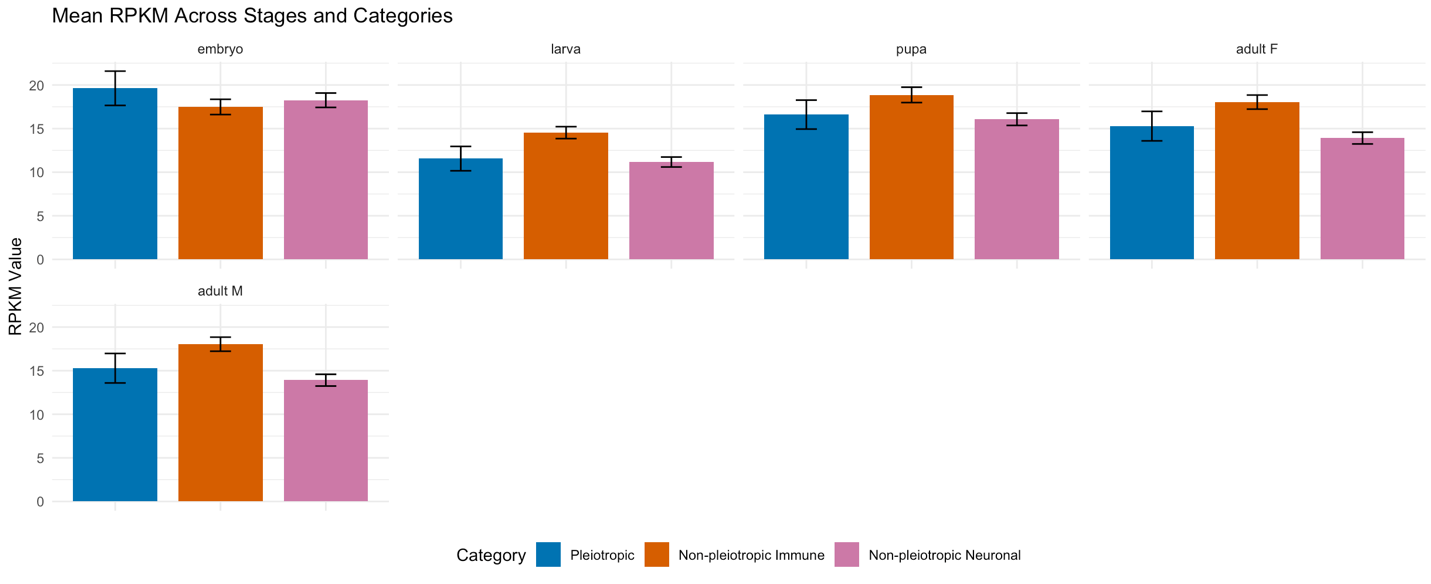

Supplemental Figures 2. Stage-specific expression distributions of genes across developmental stages, measured as RPKM values from RNA-Seq data. Violin plots depict the distribution of RPKM values across stages (embryo, larva, pupa, adult female) for three gene categories: pleiotropic genes (plei), non-pleiotropic immune genes (nonplei_immune), and non-pleiotropic neuronal genes (nonplei_neuro) (Top Panel). Bar plots show mean RPKM values for each category and stage, providing a comparison of average gene expression levels across developmental time points (Bottom Panel).

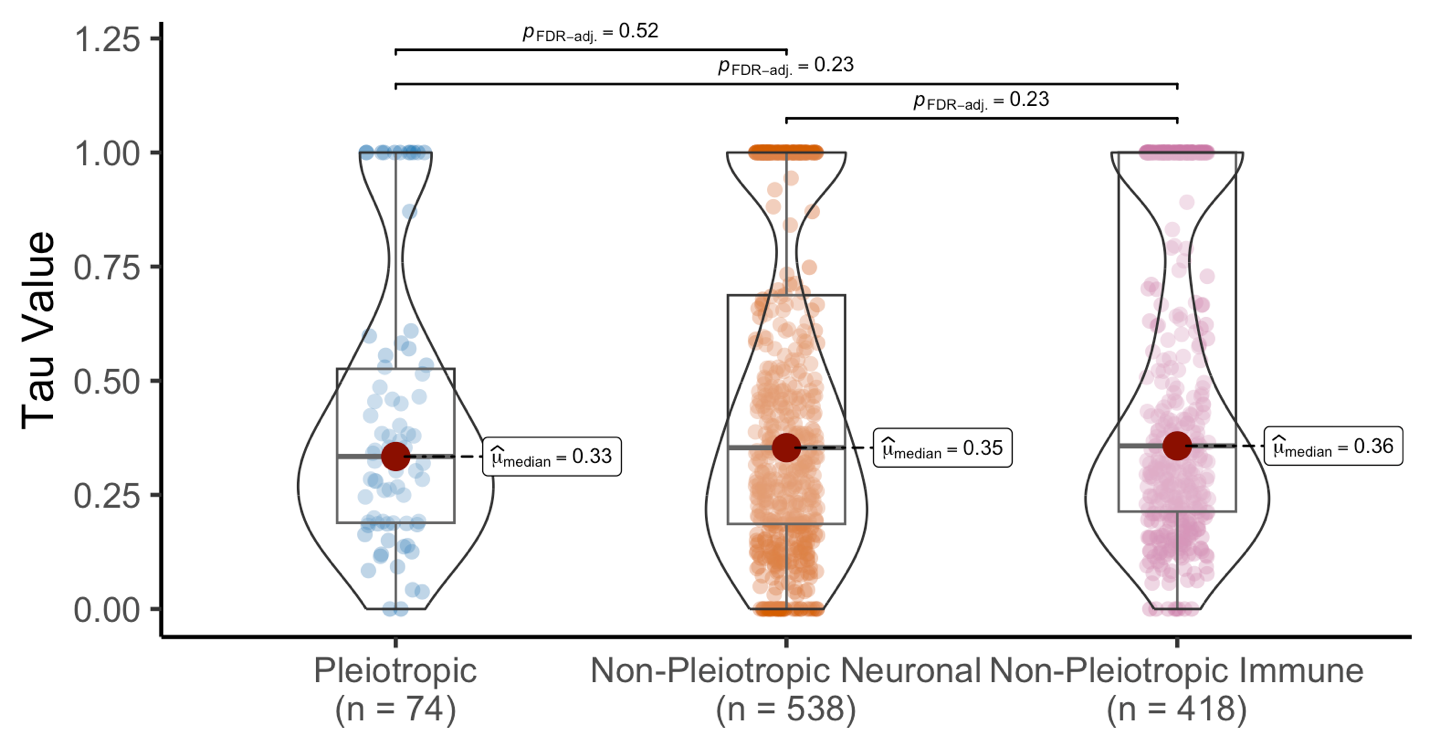

Supplemental Figure 3. Inclusion of embryonic expression increases developmental stage specificity (τ) across gene classes. Gene expression specificity (τ) was calculated across four developmental stages including embryo (6–8 hr), larva (L1), pupa (P1), and adult male (AdM1) for pleiotropic, non-pleiotropic neuronal, and non-pleiotropic immune gene classes. Tau values range from 0, indicating broad expression across stages, to 1, indicating expression restricted to a single developmental stage. Violin plots show the density distribution of τ values for each gene category, with individual points representing τ values for individual genes and overlaid boxplots indicating the median and interquartile range. Post hoc pairwise Dunn tests were used to calculate pairwise comparisons, and p values were adjusted for false discovery rate (Benjamini-Hochberg FDR).
